# Multimodal single-cell analysis uncovers transcription factor networks underlying T-cell aging

**DOI:** 10.64898/2026.05.19.726137

**Authors:** Mina Shaigan, Deepika Puri, Giulia Fornero, René Krüger, Mara Steiger, Hannes Klump, Alexander Meissner, Helene Kretzmer, Wolfgang Wagner, Ivan G. Costa

## Abstract

Aging of the immune system is associated with chronic inflammation and impaired immune function, yet the regulatory mechanisms underlying these changes remain incompletely understood. Here, we generated paired single-cell transcriptomic and chromatin accessibility profiles from peripheral blood mononuclear cells of young and old healthy donors to characterize immune aging at single-cell resolution. Using an integrative computational framework for multi-omic single-cell analysis, we detected pronounced age-associated changes in T cells, including loss of naïve CD8+ T cells and expansion of differentiated memory and effector populations. Aging was accompanied by increased inflammatory signaling and reduced oxidative phosphorylation programs. Enhancer-based gene regulatory network analyses identified a reduced role of TCF7 and increased activity of inflammatory regulators, including FOSL2, in aged T cells. Integration with genetic association and eQTL datasets further supported the functional relevance of age-associated regulatory regions and their target genes.

## 1 Introduction

Aging is a fundamental biological process that drives widespread molecular and cellular alterations, including the accumulation of cellular damage [1], diminished regenerative capacity [2], and increased disease susceptibility [3] as a consequence of immune dysfunction. Epigenetic regulation and chromatin organization are central to this process, as underscored by the broad application of DNA methylation–based aging clocks [4]. Despite their remarkable predictive accuracy, these clocks have thus far provided only limited mechanistic insight into the molecular processes that actively drive aging.

To gain a more direct understanding of age-associated regulatory changes, recent studies have profiled chromatin accessibility in peripheral blood cells using bulk chromatin accessibility assays (Assay for Transposase-Accessible Chromatin with sequencing; ATAC-seq) to uncover aging-related alterations in immune populations. Ucar and colleagues generated bulk RNA-seq and ATAC-seq data from human peripheral blood mononuclear cells (PBMCs) obtained from 77 donors across a broad age range and reported increased chromatin accessibility at myeloid lineage genes with age, whereas regions showing reduced accessibility were predominantly associated with lymphocyte-related genes [5]. Notably, they identified an age-associated decline in IL7R chromatin accessibility, which was associated with the decrease in expression of IL7R in CD8 T cells, suggesting a differentiation bias within the CD8 T cell compartment. This is supportive of the immune aging hypothesis, which indicates that loss of naïve T cells leads to a decline in immune function [6].

Subsequently, Morandini and colleagues [7] generated PBMC ATAC-seq profiles from 159 donors and confirmed similar cell-type-specific aging signatures. Building on these findings, they developed a chromatin-based aging clock that achieved a mean absolute deviation (MAD) of 5 years within their cohort. However, when applying the same clock to an expanded dataset that integrates samples from [5, 8] the MAD increased to 17 years [7]. Importantly, correcting for cell-type composition substantially improved predictive performance, reducing the MAD to 5.27 years. This demonstrates that open chromatin signatures derived from bulk measurements are strongly influenced by cellular composition, and potentially miss cell specific changes, which are crucial to understand immune aging.

Previous work suggests that age-associated immune dysfunction arises from coordinated, cell-intrinsic regulatory alterations that cannot be fully resolved using bulk measurements. In contrast, multi-modal single-cell sequencing enables simultaneous profiling of transcriptomic states and chromatin accessibility at single-cell resolution, thereby revealing true cell-intrinsic regulatory changes from shifts in cellular composition. To address this, we generate paired single-cell RNA and ATAC sequencing data from peripheral blood mononuclear cells (PBMCs) obtained from young and old healthy donors to systematically dissect the molecular basis of immunoaging. Beyond defining age-associated changes in cell-type composition and transcriptional programs, our multimodal single-cell data enables inference of cellular differentiation trajectories and reconstruction of enhancer-driven gene regulatory networks that govern immune cell identity and state transitions [9, 10, 11]. Together, these analyses provide a comprehensive, high-resolution characterization of immune aging, revealing coordinated epigenomic, transcriptional, and gene regulatory changes that drive age-associated immune decline.

## 2 Results

### 2.1 Single-cell multi-omics integration

To characterize regulatory mechanisms related to epigenetic aging in blood cells, we collected PBMCs from five healthy young adults (19-26 years) and five healthy old adults (61-74 years). Gene expression and chromatin accessibility of single-cell profiles were profiled with a 10x Multiome essay (Fig. 1a). This allowed us to detect 86,993 high-quality cells with an average of 2,492 detected genes and a TSS enrichment of 0.25 for scATAC-seq cells (Supp. Fig. 1a and 1b). The data were integrated [12] and clustered, which allowed us to identify twenty cell types with clear gene and peak markers (Supp. Fig. 1c and 1d). In detail, we detected two monocyte cells populations (CD14^+^ monocyte and CD16^+^ monocyte), three B cells populations (naïve B, intermediate B, and memory B cells), three natural killer (NK) cells populations (NK, CD56^bright^ NK, and NK proliferating), nine T cells populations (Naïve CD4^+^ T, Memory CD4^+^ T, Naïve CD8^+^ T, Central Memory CD8^+^ T, Effector Memory CD8^+^ T, *γδ* T (gdT), regulatory T (T-reg), mucosal-associated invariant T (MAIT T), double-negative T (dnT)), conventional dendritic cell (cDC), plasmacytoid dendritic dell (pDC), and intermediate cell type which genes of both NK and CD8^+^ T expressed (intermediate NK-CD8) (Fig. 1b; Supp. Fig. 1c and 1d).

**Figure 1:**
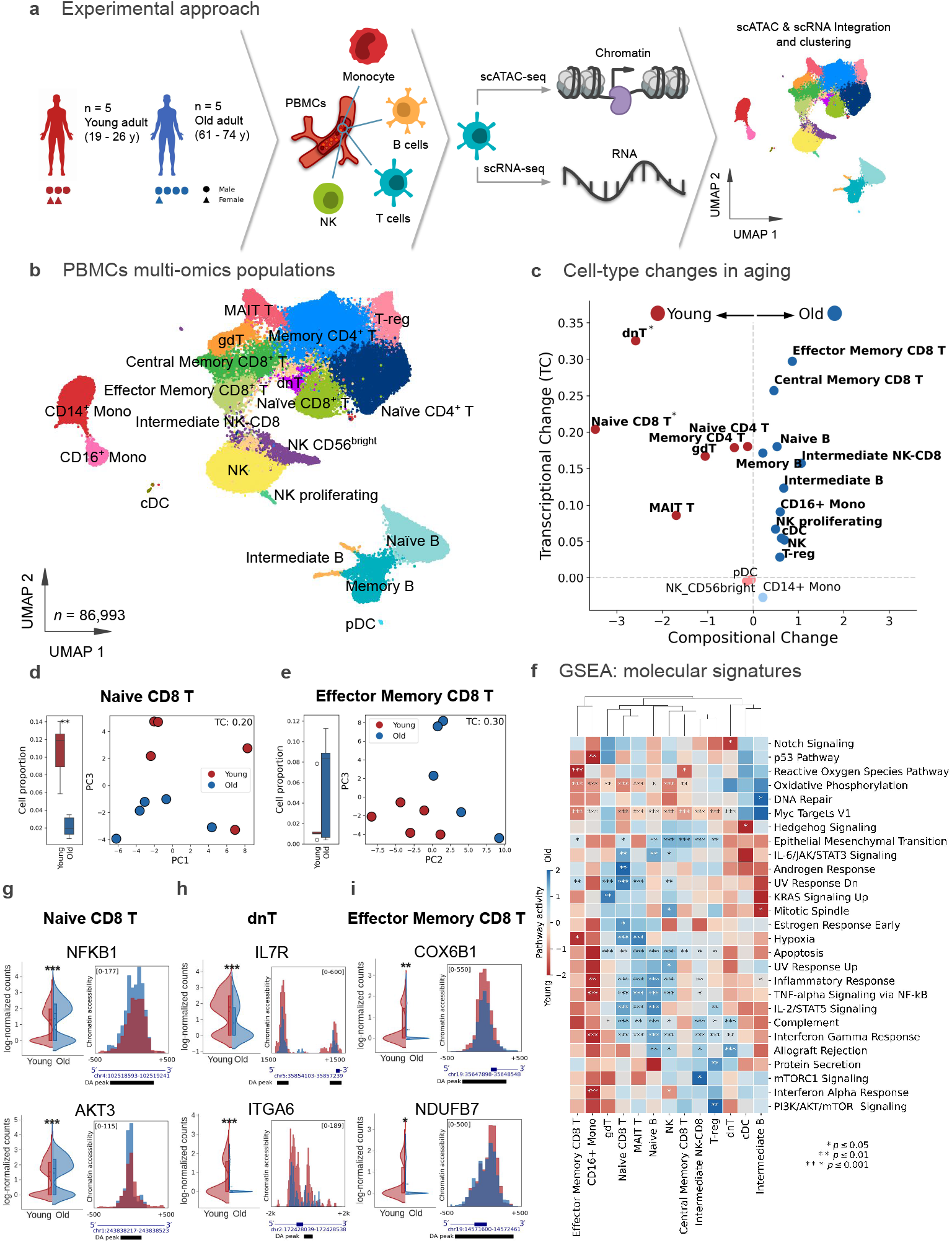
Single-cell multi-omics analysis of PBMCs. a) Schematic of single-cell multi-omics analysis of PBMCs from five healthy young adults and five healthy old adults. b) UMAP showing PBMC single cell clusters. c) A scatter plot showing Cohen’s D compositional change as the size of the difference between young and old groups means (x-axis) and transcriptional changes (y-axis) for all detected cell types. Colors correspond if the cell type is more frequent in younger (red) or older blue) donors. A Mann–Whitney U test showed that the cell proportion distributions differed significantly between young and old groups in dnT and naïve CD8^+^ T cells (p-value < 0.05). d) Boxplot showing cell proportion change for Naïve CD8^+^ T cells across young and old donors (left) and the principal component analysis of pseudo-bulk scRNA-seq (right) showing the transcriptional change between young and old donors for this cluster. Transcriptional change measures how separated old (blue color) and young (red color) samples are in the PCA space using the silhouette index of the two PCs with the highest correlation with aging. e) The same as d) for Effector Memory CD8 T cells. f) Gene Set enrichment analysis of DEG genes. We only show terms and cell types with at least one significant association with young or old DEGs. The two-sided p-values are derived from the t-test of the regression coefficient in the univariate linear model. Red color indicates enrichment in young donors, while blue color indicates enrichment in old donors.g, i) Examples of genes with differential expression and differential accessible peaks in naïve CD8^+^ (g) and Effector Memory CD8^+^ T cells (i). Red colors indicate young donors and blue colors to old donors. P-values is f-i) are represented as ^*\*\*\**^*p* < 0.001, ^*\*\**^*p* < 0.01, ^*\**^*p* < 0.05.

Next, we explored the age-related transcriptional changes in the detected cells by considering how well donor specific cell type expression can separate old and young samples (Fig. 1c to 1e; Supp. Fig. 2a; Methods Sections). We complemented this approach with an analysis of cell composition, assessing changes in the distribution of cell proportions across replicates by comparing young and old samples. Altogether, T cells were among the ones with the highest age-related expression changes. Regarding compositional changes, naïve CD8^+^ T cells and dnT cells showed a decline in cell numbers with age, whereas memory T cells (both effector and central subtypes) increased in abundance over time. We also observed a tendency of B cells to decrease with age and monocytes to increase with aging. Despite these cell composition changes, these cell types displayed less pronounced expression profiles changes than T cells (Fig. 1c). Overall, these results indicate that immune cell composition shifts toward an effector and inflammatory state of T cells with age, associated with a loss of progenitor T cells in line with previous studies [5].

**Figure 2:**
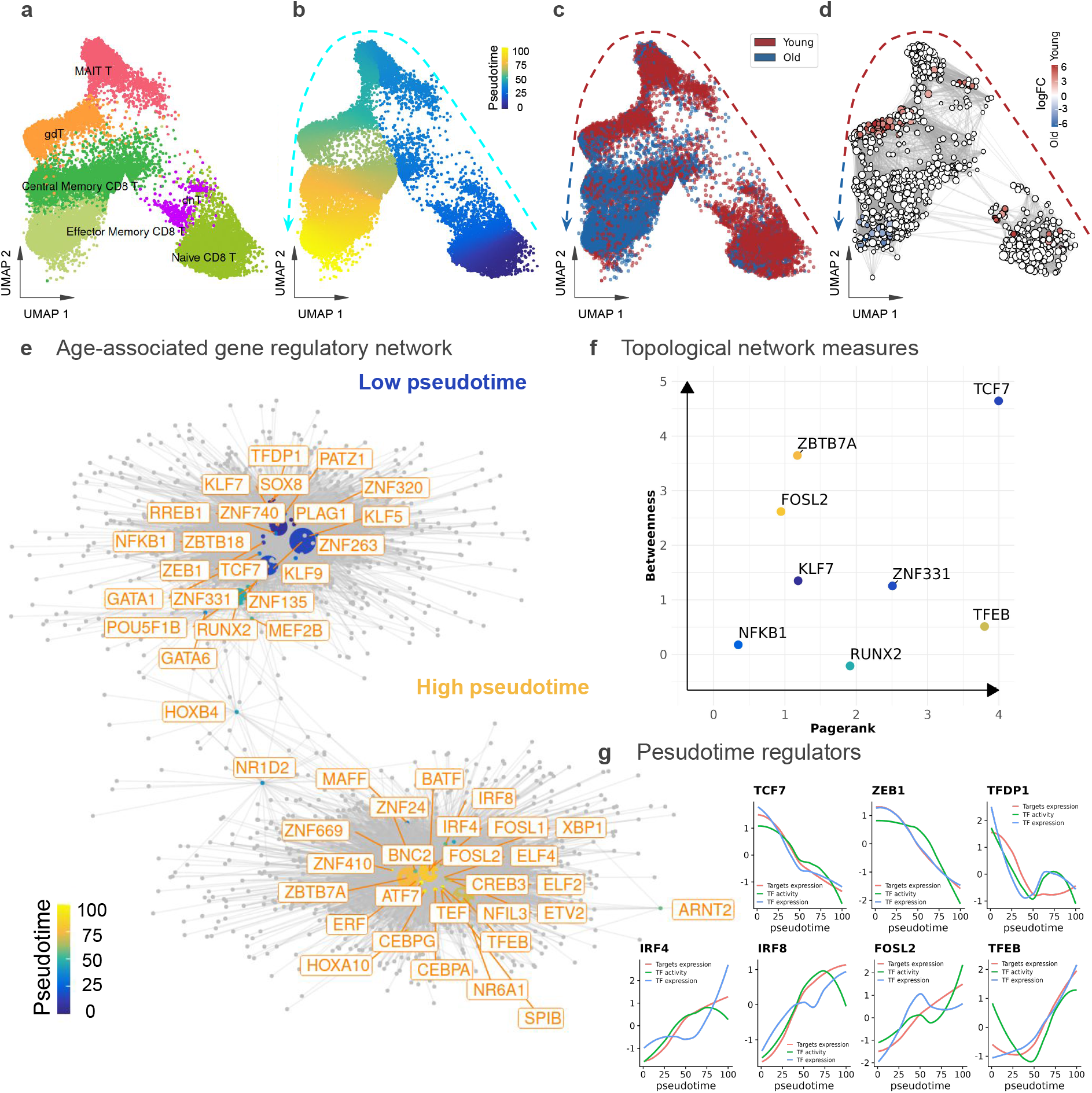
Gene regulatory network in aging. a,b) UMAP showing cell types (a), pseudotime estimates (b) and age group of donor (c). d) Milo based cell neighborhoods representation. Color nodes represent cells neighborhoods with differential abundance (FDR 10%). Young neighborhoods are shown in red and old neighborhoods are shown in blue. e) Visualization of the Inferred GRN for age-related T-cell. Each node represents a regulator TF (orange labels) or a target gene (gray dots). Each TF is colored based on their average pseudo-time estimates. The size of the TFs is proportional to network importance statistics (page-rank). f) Scatterplot showing top TFs regarding eGRN network betweenness (y-axis) and pagerank (x-axis) measures. Higher values indicate higher importance as regulators. Colors represent the TF pseudo-time. g) Line plots showing the TF binding activity (green), TF expression (blue) and average target expression (red) (y-axis) along the age-related T-cell trajectory (x-axis) for selected TFs.

Next, we performed GSEA (Gene Set Enrichment Analysis) using KEGG pathways to evaluate the functional relevance of the cell type specific changes. We observed an increase in inflammatory signaling over aging in most cells, but also a decrease in oxidative phosphorylation in aging (Fig. 1f). In naïve CD8^+^ T cells, for example, we observe the gain of chromatin accessibility and expression of NFK*β* and the longevity regulating pathway related gene Akt3 (Fig. 1g). While for dnT cells, we detected the loss of accessibility and expression in IL7R as part of JAK-STAT signaling and PI3K-AKT signaling pathways, and the IL-2/STAT5 signaling related gene ITGA6 (Fig. 1h). Of note, the IL7R peak is the same as discussed in a bulk based analysis [5], but our analysis can show now that the peak changes are mostly related to early T cells (dnT but also naïve CD8^+^ T cells), as an example of the importance of single cell analysis (Supp. Fig. 2c). For Effector Memory T CD8^+^ T cells, we also see a decrease of gene expression and accessibility of COX6B1 and NDUFB7 genes, which are both related to oxidative phosphorylation (Fig. 1i).

Altogether, these results support the fact that our multimodal data can characterize transcriptional, compositional, and chromatin changes through age. Remarkably, we observed that these changes support a transcriptional rewiring of T-cells, which acquire inflammatory and oxidative stress programs. Moreover, an age specific loss of naïve T cells supports an imbalance in T-cell differentiation in aging.

### 2.2 T-cell gene regulatory network

Due to the prominent immune aging phenotype of T cells, we next characterized gene regulatory networks with a focus on the T cell compartment and related this to aging. We first performed a cell trajectory and pseudo time analysis using ArchR [13] (Fig. 2a and 2b) . The trajectory indicates naïve CD8^+^ T cells in the beginning of the trajectory (low pseudotime) and Effector Memory CD8^+^ T cells in the end of the trajectory (high pseudotime). Moreover, cells from young donors tend to have low pseudo time and cells of old donors to have high pseudotime, which is supported by Milo based abundance testing (Fig. 2c and 2d; Supp. Fig 3a). These results suggest that the trajectory, associated with cell differentiation, captures a shift from naïve to effector T cells during PBMC aging.

**Figure 3:**
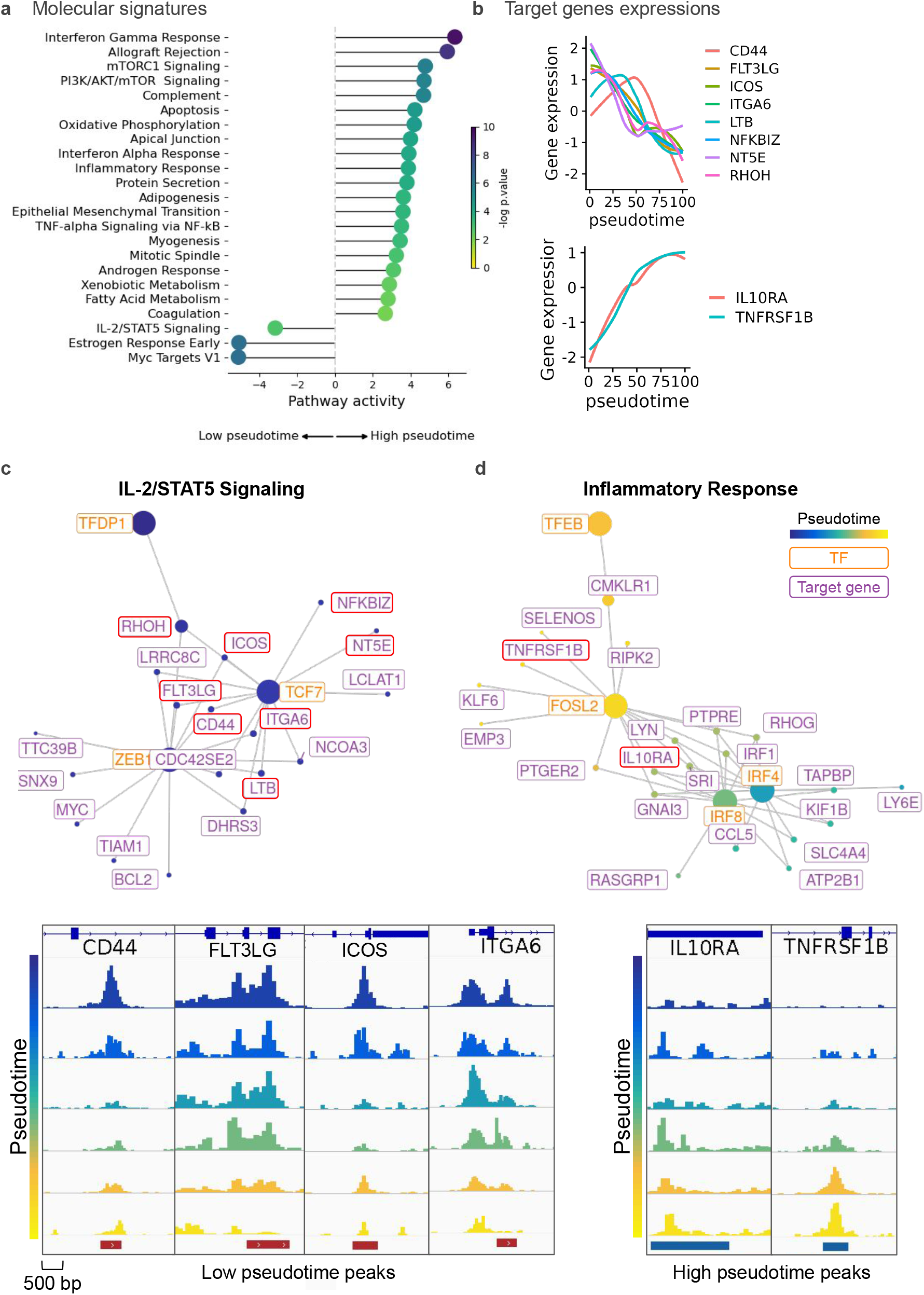
Pathway-specific Gene regulatory network. a) Gene set enrichment analysis of pathways in the eGRN modules (high vs. log pseudo-time). We compared pathway activities in target genes regulated by low pseudotime and high pseudotime TFs using an univariate linear model. We considered the average correlation between TF activity and gene expression as the regulator weight for each gene. The bar plot shows the significant signatures (p-value < 0.01) with color coding based on the negative log p-value. b) We showed the expression of selected target genes (y-axis) along the pseudotime trajectory (x-axis) associated with IL-2/STAT5 and inflammatory pathways. c) eGRN associated with IL-2/STAT5 signaling pathway and chromatin changes along pseudotime for selected target genes. d). Same as before for the eGRN related to the inflammatory response pathway.

We next estimate an enhancer based regulatory networks (eGRN), as this will allow us to characterize transcription factors controlling the cell differentiation events. TFs are particularly important because they control age-related changes in T cells leading to rewiring of underlying regulatory programs. This eGRN is constructed by considering both the expression and activity of TFs, the binding of these TFs to enhancer regions of genes and the expression of genes using scMEGA [9]. scMEGA predicted a GRN network with a total of 52 transcription factors and 1,939 target genes (Fig. 2e). It contains two main modules, one associated with low-pseudotime (young group) cells and another with cells enriched from high pseudo time (old group) (Fig. 2e).

To delineate important TFs, we explored network properties such as TF betweeness and pagerank [14], which indicates TFs regulating other highly connected TFs and many target genes (Fig. 2f.) One particular TF, which achieves the highest values in both metrics in TCF7, which is part of the WNT signaling pathway and related to differentiation of naïve T cells [15]. An inspection of the TCF7 activity expression and target gene expression indicates a clear decrease of the values in pseudo-time. Moreover, age related TC activity decrease is particularly high in naïve CD8+ T cells (Supp. Fig. 4). In addition, we observed ZEB2, which is also involved in T cell immune function [16], and TFDP1, which is related to the cell cycle, exhibiting decreases in both pseudo-time. FOS2L, which was also highlighted by our network statistics, is part of the AP1 family and has been related to the acquisition of inflammation phenotypes by T cells [17]. This TF has an increase in activity in older donors in memory CD8 T cells (Supp. Fig. 4). Another TF selected in our network is the stress response related TF TFEB [18], which shown both pseudo-time and age related TF activity increase (Fig. 2g and Supp. Fig. 4). We observed altogether 27 TFs with increasing activity with age (Fig. 2c), notably the interferon related IRF8 and IRF4 associated with the activation of T helper cells [19, 20].

**Figure 4:**
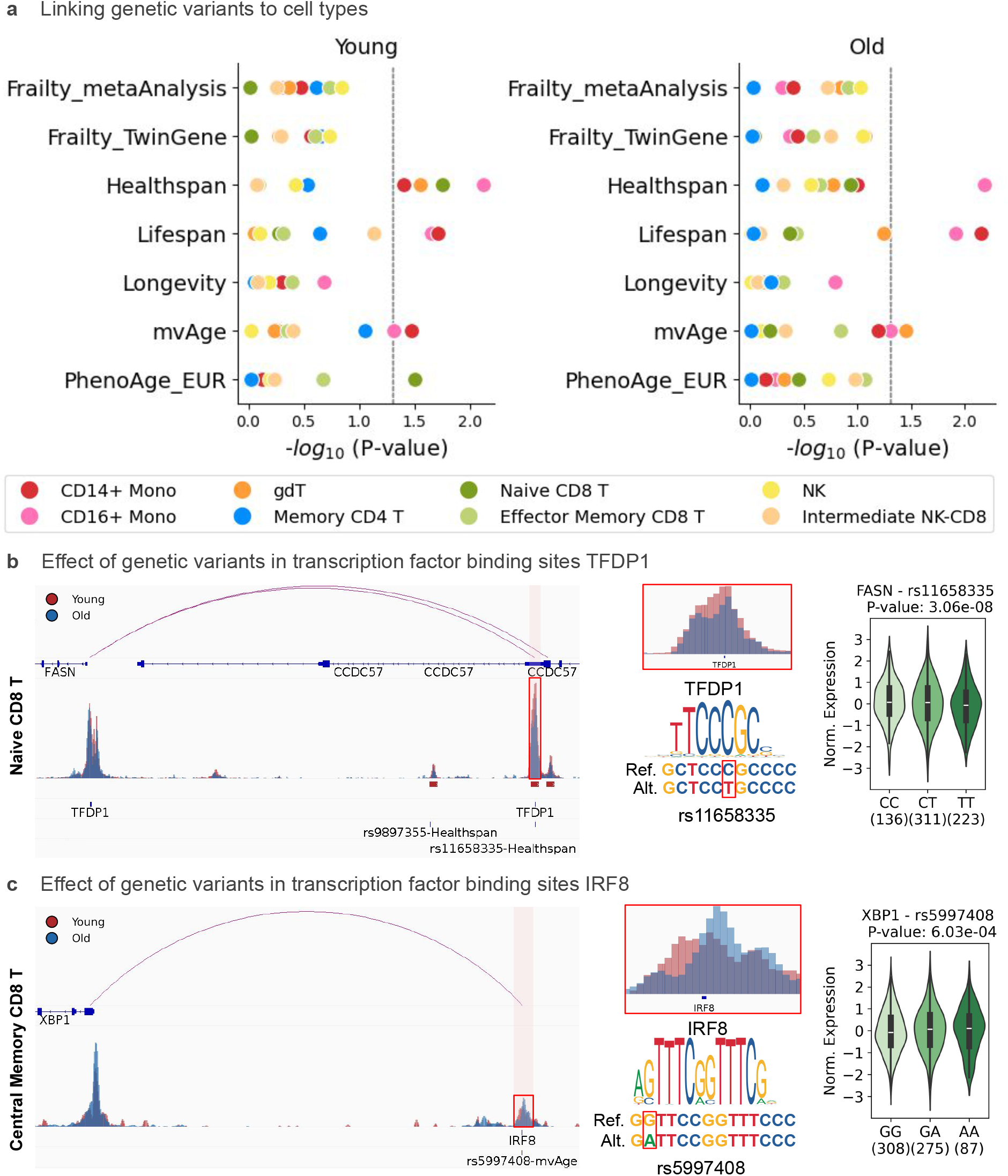
Genetic variants effects in aging. a) Cell-type specific LDSC analysis to test whether genetic variants associated with a trait are enriched in significant differential single-cell peaks in particular cell types.b, c) Chromatin accessibility (left) around the FASN and XBP1 genes, which include enhancers with transcription factors binding sites (TFDP1 and IRF8) disrupted by aging related SNPs in naïve CD8^+^ T and central memory CD8^+^ T. Expression quantitative trait loci (right) for a given pair of corresponding gene and variant based on Genotype-Tissue Expression (GTEx) whole blood dataset.

### 2.3 Pathway changes in T cell aging

To further characterize molecular mechanisms associated with these eGRN modules, we performed a GSEA analysis of TFs and targets from each module (Fig. 3a). We observe a decrease in IL2-STAT5 signaling in low-pseudo time cells (Fig. 3c), while an increase in inflammatory response (Fig. 3d), apoptosis and oxidative phosphorylation is observed in high-pseudotime cells. These pathways are in accordance with an inflammation and senescence phenotype of T cells in aging [21].

Several genes of interest emerged from the pathway-specific GRN analysis, which all display a concordant increase or decrease in expression in pseudo-time, as supported in the pathway analysis (Fig. 3b). In the IL-2/STAT5 signaling network, we found that several genes were highly expressed in naïve CD8^+^ T cells and were regulated by the TCF7 and ZEB1 TFs. For example, CD44, a molecule expressed on T cells, plays a crucial role in T cell activation, homing, and migration [22]. The Inducible T-cell Co-Stimulator (ICOS) acts as a key co-stimulatory receptor on activated T cells, enhancing their proliferation and cytokine secretion [23], while ITGA6 (Integrin alpha-6) contributes to T cell adhesion and migration within tissues [24]. All these genes shown both a decrease in accessibility and expression along pseudo-time (Fig. 3b and 3c), which support the loss of their function in age.

In contrast, within the inflammatory response network, we observed increased expression of certain genes along the pseudotime trajectory. Notably, IL10RA, which is significantly regulated by FOSL2, IRF4, and IRF8 TFs, which is associated with T cell exhaustion [25], and TNFRSF1B (also known as TNFR2), is regulated by FOSL2 and XBP1 TFs, and is associated with inflammatory phenotypes of t cells [26] and has been identified as a marker for exhausted CD8+ T cells in cancers [27]. These results show that our analysis can not only predict TFs associated with T cell aging, but also establish functional connection between these TFs and crucial changes in age-associated immune function including T cell exhaustion, chronic inflammation and senescence.

### 2.4 Genetic variant effects in aging

Genome-wide association studies (GWASs) have revealed numerous genetic loci linked to age-associated traits [28, 29, 30, 31, 32, 33] offering valuable insights into the functional effects of risk variants that are enriched in chromatin regions with cell type and age specific accessibility. Since most GWAS traits are in non-coding regions and potentially affect transcription factor binding sites, they provide a unique resource to validate eGRN predictions [34, 35]. We therefore resort to genome-wide association studies to verify whether age-associated SNPs potentially affect these binding sites; and to expression Quantitative Trait mappings to validate the effect of the SNPs in the downstream genes.

For this, we performed a Linkage Disequilibrium Score Regression [36] to relate traits from GWAS from distinct studies with cell type and age specific OC regions (Fig. 4a). GWAS studies include ones related to increasing life span (mvAge [28], healthspan [29], parental lifespan [30], and exceptional longevity [31]), or aging traits (Fraility [32] and age acceleration [33]).

Notably, we observe that monocyte peaks (gain or lost in age) are associated with positive disease traits implying their central role in immune surveillance and inflammation regulation [28]. Regarding T cells, peaks specific to young donors in naïve CD8^+^ and gdT cells are both associated with health-span associated traits, which indicate that the loss of functionality of these cells represents an aging hallmark. An interesting hit is the SNP (rs11658335) associated with healthy span and disrupts a TFDP1 binding site in a naïve CD8^+^ T cell enhancer linked to Fatty acid synthase (FASN). FASN has been previously shown to be associated with cell senescence [37] (Fig. 4b). We next resort to expression quantitative trait loci (eQTL) associations from the Genotype-Tissue Expression (GTEx) whole blood dataset [38]. This indicates that rs11658335 is significantly impact on the expression of the FASN gene (cis-eQTL *P* = 3.06 *×* 10^*−*8^), with the C allele linked to higher expression levels and the T allele linked to lower expression, which again corroborates to a disruption of the TFBP1 binding site leading to a lower expression of FASN.

Another interesting hit is a SNP (rs5997408) that disrupts an IRF8 binding site on an old age related Central Memory CD8 T cell peak. This peak is associated with the X-box-binding protein-1 (XBP1), which plays a key role in regulating endoplasmic reticulum stress and T cell metabolic function (Fig. 4c). Additionally, we observed a cis-eQTL for XBP1 associated with rs5997408 (*P* = 6.03 *×* 10^*−*4^), supporting the effect of the G to A allele change on gene expression, which further support our eGRN prediction. These (and additional examples Supp. Fig. 5a and 5b) support the functional relevance of TF-enhancer links predicted in our T cell aging gene regulatory network.

## 3 Discussion

Here, we present a multimodal single-cell analysis of human immune aging that integrates transcriptional, chromatin accessibility of PBMCs from young and old donors. Our results provide a high-resolution view of how aging reshapes immune cell identity, with a particular emphasis on T cell–intrinsic regulatory changes. A main finding is that aging is accompanied by cell-type–specific transcriptional and epigenetic changes most prominently within T cells. Consistent with prior studies [5, 39], we observe a decline in naïve CD8^+^ T cells and a shift toward more differentiated memory and effector states, reflecting a loss of immune plasticity. Importantly, our data supports the value of single cell based analysis to disentangle cell specific from age specific changes. At the molecular level, aging is associated with a transition toward pro-inflammatory and stress-related programs, alongside reduced metabolic activity. These changes are reflected in both gene expression and chromatin accessibility, including decreased IL7R signaling and increased NF-*κ*B–associated activity. Together, these findings support the concept of “inflammaging” and suggest a broad reprogramming of immune cell function rather than isolated gene-level effects.

Trajectory analysis further revealed a continuous transition from naïve to effector-like T-cell states that correlated with donor age. This supports the notion that aging promotes progressive differentiation and loss of T-cell plasticity. Reconstruction of enhancer-based gene regulatory networks identified transcription factors orchestrating these transitions. Among these, TCF7 emerged as a central regulator associated with naïve T cells and with decrease of functionality in aging. TCF7 is a well-established regulator of T-cell stemness and memory potential [40], and its reduced activity in aged naïve CD8+ T cells is consistent with previous reports linking loss of TCF7 function to T-cell exhaustion and impaired immune responses [41]. In contrast, transcription factors associated with inflammatory and stress-related responses, including FOSL2, IRF4, and IRF8, showed increased activity in aged cells, suggesting a shift toward activated and dysfunctional regulatory programs during aging.

Aging of the adaptive immune system is closely tied to dysregulation of homeostatic cytokine networks. Our data showing reduced chromatin accessibility and expression of IL-2/STAT5–associated regulatory programs in aged T cells suggests an impairment of activation and survival signaling, which likely contributes to the diminished proliferative capacity and altered differentiation trajectories characteristic of immune aging. Together, these changes reflect a coordinated decline in cytokine-mediated regulatory circuits that underpin both homeostatic maintenance (IL-7) and activation competence (IL-2) of T cells, consistent with broader hallmarks of immunosenescence [42, 43].

Some limitations should be noted. The relatively small size in the single-cell cohort limits statistical power and investigation of sex specific change. While these associations provide a functional hypothesis connecting genetics to regulatory aging, they remain correlative and require experimental validation to establish causality. In summary, our findings support a model in which immune aging is driven by coordinated, cell-type–specific regulatory changes, particularly in T cells, and support the importance of multimodal single-cell approaches for dissecting intrinsic molecular changes from cell compositional effects. This paves the way for future studies incorporating larger, more diverse cohorts and functional perturbations will be essential to establish causality and identify regulators of immune aging.

## 4 Methods

### 4.1 Blood sample collection and PBMC isolation

Peripheral blood samples were collected from healthy adults between the ages of 20 and 70, each of whom provided informed written consent to participate. The study followed the ethical principles of the Declaration of Helsinki and received approval from the RWTH Aachen University ethics committees (EK 206/09 and EK 041/15). After collection, whole blood was placed into EDTA-coated vacutainer tubes and kept at 4°C for few hours before further processing.

Peripheral blood mononuclear cells (PBMCs) were isolated from the collected samples using density gradient centrifugation with Pancoll (Pan-Biotech). In summary, each blood sample was mixed with an equal volume of phosphate-buffered saline (PBS), then carefully layered onto Pancoll and centrifuged as recommended by the manufacturer. The PBMC layer was gently collected and transferred to a fresh 50mL conical tube. To ensure purity, cells were washed two to three times with PBS to remove platelets and any remaining plasma. Finally, the cell pellets were resuspended in PBS or culture medium, and the number of viable PBMCs was determined using trypan blue exclusion with either a hemocytometer or an automated cell counter.

### 4.2 Single-cell multi-omics experiments

Peripheral blood mononuclear cells (PBMCs) samples were obtained from five young (19-26 years) and five old (61-74 years) healthy human adults and were further used to create a single-cell multi-omics library. PBMCs freshly isolated from whole blood of healthy donors using the Chromium Next GEM Single Cell Multiome ATAC + Gene Expression (GEX) protocol (CG000338). For the Nuclei Isolation step the protocol CG000365 was followed.

Granulocytes were removed by freezing the samples in DMSO-containing medium after PBMC isolation, followed by thawing and centrifugation-based cleanup prior to library preparation in Cologne Centre for Genomics (CCG).

The single-cell RNA sequencing and single-cell Assay for Transposase Accessible Chromatin using sequencing (ATAC-seq) samples were processed with the 10X Multiome kit, scATAC libraries were sequenced 51-8-24-51, scRNA libraries were sequenced 29-10-10-89, both on NovaSeq 6000.

### 4.3 Single-cell ATAC-seq data quality control

The single-cell data were pre-processed using Cell Ranger Arc 2.0.1 with GRCh38 as a reference genome. In short, raw Illumina binary base call (BCL) files were converted into FASTQ files for both ATAC and GEX datasets using cellranger-arc mkfastq. Then, ATAC reads and GEX reads are aligned to the GRCh38 (hg38) human reference sequence using STAR. Barcodes and molecular features were filtered and counted.

Using the ArchR package [13], the fragment files were converted to Arrow files. The low-quality cells are filtered out using minTSS = 15, minFrags = 1,000, and maxFrags = 100,000 to retain cells with TSS enrichment score higher than four and number of unique fragments between 1,000 and 100,000. The detected doublets were removed using ArchR. A Tile Matrix, which defines a fixed-width (500 bp) sliding window of bins across the whole genome, was used for dimension reduction (using iterative LSI with 30 dimensions) and clustering. Moreover, the Harmony batch-corrected reduced dimensions handled technical variations or batch differences among samples.

### 4.4 Single-cell RNA-seq data quality control

The downstream analyses for the scRNA-seq dataset were performed using Seurat [44]. We filtered cells that have unique feature counts less than 200 or over 2,500, or higher than 20 percent mitochondrial counts. We normalized the data using the LogNormalize method, which adjusts the feature expression measurements for each cell based on the total expression, then multiplies this by a scale factor of 10,000, and finally applies a log transformation to the result. Considering 5,000 genes with high cell-to-cell variation for each sample, we chose 3,000 genes to integrate the samples. To do so, sample datasets were projected into a shared low-dimensional space using Reciprocal PCA with 50 dimensions. In this way, Seurat could find anchors as reference points to understand how datasets related and matched cells that are closest while correcting for batch effects.

### 4.5 scATAC-seq and scRNA-seq integration and clustering

We used the previous dimension reduction spaces (reciprocal PCA for scRNA-seq and LSI for scATAC-seq) as input for MOJITOO to find a shared space of multimodal RNA and ATAC cells, which is based on 30 canonical components. This was given as input for a Louvain algorithm with resolution 0.5, which obtained 23 clusters of 86,993 valid cells. Next, the Uniform Manifold Approximation and Projection (UMAP) ran on this reduced dimensional space for visualization purposes.

To annotate cells, we identified marker genes for each cluster using the Wilcoxon rank-sum test to compare the expression of each gene in a specific cluster against all other cells. To determine if a gene is statistically significantly expressed in each cluster, at least 25% of the cells must express the gene, and the minimum log fold change should be 0.25. Finally, we annotated clusters 0 and 15 as naïve CD4^+^ T (FHIT, CASC15, DACH1), 1 as memory CD4^+^ T (ITGB1, SEMA5A, MYO16), 2 as natural killer (GNLY, LINGO2, GRIK4), 3 as naïve B (IL4R, AL139020.1, SCN3A), 4 as central memory CD8^+^ T (CD8A, MSC-AS1, TPRG1), 5 as naïve CD8^+^ T (CD8B, NRCAM, ROBO1), 6 as effector memory CD8^+^ T (TRGC2, LINC02086, KLHL4), 7 as CD14^+^ monocyte (IL1B, VCAN-AS1, TNFAIP6), 8 as memory B (COCH, SSPN, COL4A3), 9 as gamma delta T (rna TRGC1, TRDC, SIPA1L2), 10 as regulatory T (CTLA4, RBMS3, NEBL), 11 as mucosal-associated invariant T (SLC4A10, AC010967.1, ME1), 12 as CD56^bright^ natural killer (PPP1R9A, KIT, ZMAT4), 13 as CD16^+^ monocyte (CDKN1C, MEG3, C1QA), 14 as intermediate natural killer and CD8^+^ (GNLY, PPP1R9A, STXBP6), 16 and 18 as intermediate B (MS4A1, LINC02227, SOX5-AS1), 17 and 19 as double negative T (MIR4422HG, MYB, ZNF704), 20 as proliferating natural killer (MKI67, E2F7, DIAPH3), 21 as conventional dendritic cell (FLT3, CYYR1, LINC01478), and 22 as plasmacytoid dendritic cell (MZB1, IGHG1, IGKC). We doubled check markers with ones defined in the Azimuth human PBMC reference [45].

To have more robust cell specific peaks, we created pseudo-bulk tracks for every cell type and performed peak calling using ArchR (addReproduciblePeakSet function with default setting). Peaks merged to get the consensus peak set, resulting in 233,399 peaks. We identified differential accessibility peaks using the Wilcoxon rank-sum test on the matched cells of cell types. Peaks filtered by log fold change less than 0.01 and FDR higher than 0.05.

### 4.6 Identification of age related molecular and cellular features

We considered the 21,923 genes and filtered out genes detected in fewer than three cells. We normalized each cell by total counts over all genes using CPM (Counts Per Million) and log-transformed the result. After keeping genes that are expressed in more than 20% of all cells, we next used MAST [46] to perform differential expression analysis for each cell type in aging. MAST finds genes within cells that are expressed differently between the young and old groups (as fixed effects), while accounting for biological variability between replicates (as random effects) using a generalized linear mixed effects regression model. If genes have an absolute log fold change value higher than 1 and a false discovery rate (FDR) lower than 0.05, they would be considered significantly differentially expressed.

A total of 233,399 peaks were CPM-normalized per cell and log-transformed. We filtered for peaks accessible in at least three cells. Using MAST, we identified peaks that were differentially accessible (DA) between young and old samples. Peaks with an absolute log fold change greater than 0.01 and a false discovery rate (FDR) below 0.05 were considered significantly differentially accessible.

We used Enrichr [47, 48, 49] from GSEApy [50] to obtain gene sets from the Human Molecular Signatures Database (MSigDB) [51, 52] and the aging atlas database [53]. Given the direction and size of changes in each gene, we applied a univariate linear model (ULM) (Decoupler [54]) to understand which pathways best explain changes associated with aging across cell types.

To analyze cellular composition, we computed each sample’s proportion of cell types. Then, using the two-sided Mann-Whitney U-test (MWU), we compared the distributions of cell sample-level frequencies between the young and old groups and whether there was a significant difference between groups. To measure how large the difference is between the groups, we used Cohen’s D Score since there were only five samples per group so that it would be more useful for interpretation.

Finally, to access the transcriptional change in aging, we combined principal component analysis (PCA) silhouette scoring similar to that proposed in [55]. In short, we performed pseudo-bulk for every donor and cell types and used this as input to PCA and considered PCs regarding 75% of explained variance. We then considered the silhouette index of the two PCs with the highest correlation with aging. That is, this indicates whether for this particular cell type, it is possible to distinguish between old and young donors using one of the main PCs. Examples are shown in Supplementary Fig. 2a.

### 4.7 Gene regulatory network in aging

To uncover the gene regulatory mechanisms in our data set, we performed a trajectory analysis and gene regulatory inference as described bellow. We focus here on T cells, due to their age related changes as revealed in our DEG and compositional analysis. More exactly, we only consider the naïve CD8^+^ T, dnT, MAIT T, gdT, central memory CD8^+^ T, and effector memory CD8^+^ T. We next used ArchR to estimate a trajectory on the first two CCA (MOJITOO) dimensions, where dnT cells were indicated as root cells. To further test the significant relationship between age and the pseudotime trajectory, we used the Milo abundance analysis method [56], which detects cellular neighborhoods enriched in a particular condition (old or young).

We next used scMEGA to find an enhancer based Gene regulatory network. scMEGA first identifies functionally active transcription factors (TFs), whose TF activity scores and gene expression are correlated (R > 0.4; p-value < 0.01) and show trajectory-specific changes. TF activity was estimated with chromVAR[57] using JASPAR version 2022 motifs [58]. To select relevant target genes, scMEGA first considered the top 10% variable genes across the T cell pseudotime trajectory. Then, the genes linked to peaks based on the correlation between gene expression from the scRNA-seq and peak accessibility from scATAC-seq data along the T cell pseudotime trajectory (adjusted p-value <1e-04). In this way, only potential target genes are selected that are associated with at least one peak (that can be seen as a cis-regulatory element like an enhancer), which is at least 2k base pairs(bp) from the transcription start site of a gene. We then connect TFs to genes whenever we find a binding site in a linked enhancer. The correlation between selected TF and target gene expression was used to build a quantitative gene regulatory network (GRN). The GRN is a directed graph from TF to gene, weighted by correlation. TF-gene links with correlations above 0.5 were retained.

With the eGRN network, we can rank nodes (in particular TFS) by using network properties such as page-rank (how influential the TF is in the network) or betweenness (how many interactions are mediated by this TF [14]. Moreover, we use a network layout algorithm (Fruchterman–Reingold) to visualize this network, and detect two gene modules (high vs. low pseudo-time genes and targets). As before, we use ULM for gene set enrichment analysis of the network modules.

### 4.8 Association of GWAS traits with age-related cell types

Considering age-cell-type specific chromatin accessibility markers peaks (with a false discovery rate (FDR) threshold of 0.05 and a minimum log fold change of 0.01 using the Wilcoxon rank-sum pair-wise test), we performed linkage disequilibrium score regression (LDSR) analysis [36, 59, 60] to relate these to Genome-wide association studies (GWAS) of human aging traits. For this, we considered traits related to healthspan [29], parental lifespan [30], exceptional longevity [31], epigenetic age acceleration (EAA) (from the PhenoAge cohort) [33], and frailty (from a meta-analysis of UKB and TwinGene cohort participants) [32]. Additionally, a multivariate GWAS incorporating these traits was performed by mvAge [28]. Traits such as mvAge, extreme longevity, healthspan, and parental lifespan are positively correlated with age, indicating slower biological aging and longer survival. In contrast, the PhenoAge trait, which estimates biological age, and the frailty trait, which measures physiological vulnerability, serve as indicators of accelerated aging. Furthermore, we overlapped transcription factor binding sites (from our eGNR analysis) in DA peaks with SNP locations for each trait to narrow the search space.

The Genotype Tissue Expression (GTEx) version 8 dataset, which comprises 670 whole blood samples, enabled us to investigate the impact of genetic variation on gene expression in cis through genotypes. Gene expression values for the samples were normalized by first selecting genes with expression above 0.1 transcripts per million (TPM) and with a minimum of 6 reads in at least 20% of the samples. Then, expression values were normalized using the trimmed mean of M-values (TMM) of the edgeR approach [38].

## 5 Acknowledgements

We thank the Cologne Centre for Genomics (CCG) for sequencing experiments. This work was funded by the German Research Foundation by grants (ID: 388802535; 458369830; SFB 1506-1/450627322).

## 6 Ethics declarations

This study was conducted in accordance with guidelines approved by the local ethics committee of RWTH Aachen University (EK 041/15).

## 7 Data Availability

Pre-processed data will be available in Zenodo (https://doi.org/10.5281/zenodo.20117453), and raw data will be deposited in EGA upon acceptance.

## 8 Author contributions

Mina Shaigan and Ivan G. Costa have designed the study, analyzed data, and revised the manuscript. Mina Shaigan performed all programming tasks and wrote the first version of the manuscript. Deepika Puri supported the analysis of the single-cell data, and René Krüger supported the GWAS analysis. Giulia Fornero has performed all wet-lab work. Ivan G. Costa, Alexander Meissner, and Wolfgang Wagner conceptualized the study and performed funding acquisition. All authors have revised and approved the final version of the manuscript.

## 9 Conflicts of Interest

W.W. is cofounder of Cygenia GmbH (www.cygenia.com) that can provide service for epigenetic analysis to other scientists. RWTH Aachen Medical School holds a patent on DNAm analysis of TCF7 for quality control of T-cells and W.W. is named as an inventor.

## Supplementary Materials

**Supplementary Figure 1:**
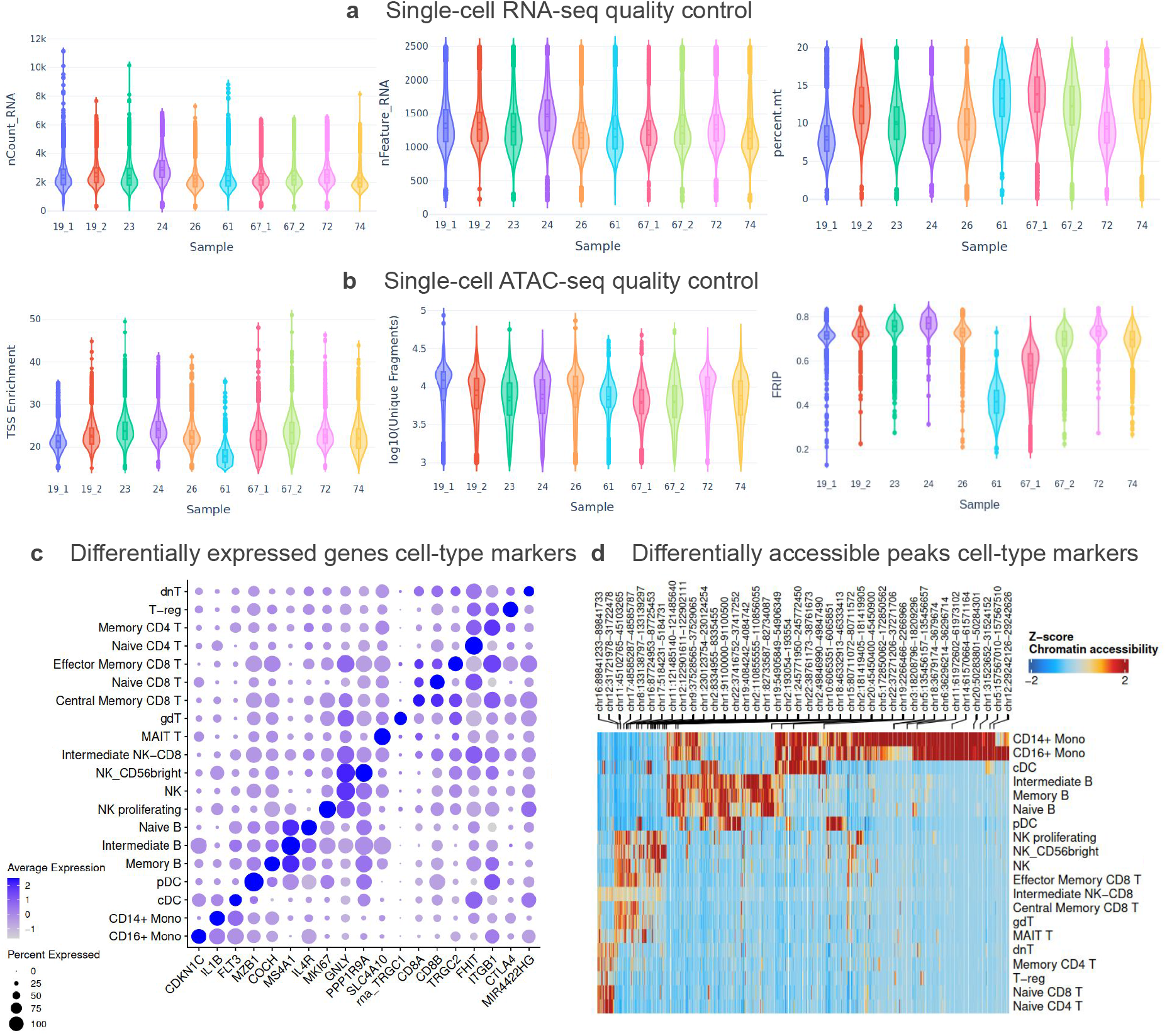
Single-cell multi-omics quality control. a) Distributions of the total UMI counts, number of unique genes, and percentage of transcripts mapped to mitochondrial genes for all single-cell RNA-seq libraries. b) Distributions of the ratio of transposition events centered on transcription start sites (TSS) compared to flanking background regions and the number of unique, non-duplicated DNA fragments mapped to the genome in single-cell ATAC-seq. c) Dot plot showing cell type specific marker genes. The size of the dot shows the percentage of cells within a cell-type, and the color indicates the average expression level of all cells within a cell-type. d) Differential accessibility peaks across cell-types filtered with FDR <= 0.1 and Log2FC >= 0.5. Peak accessibility is standardized using z-scores.

**Supplementary Figure 2:**
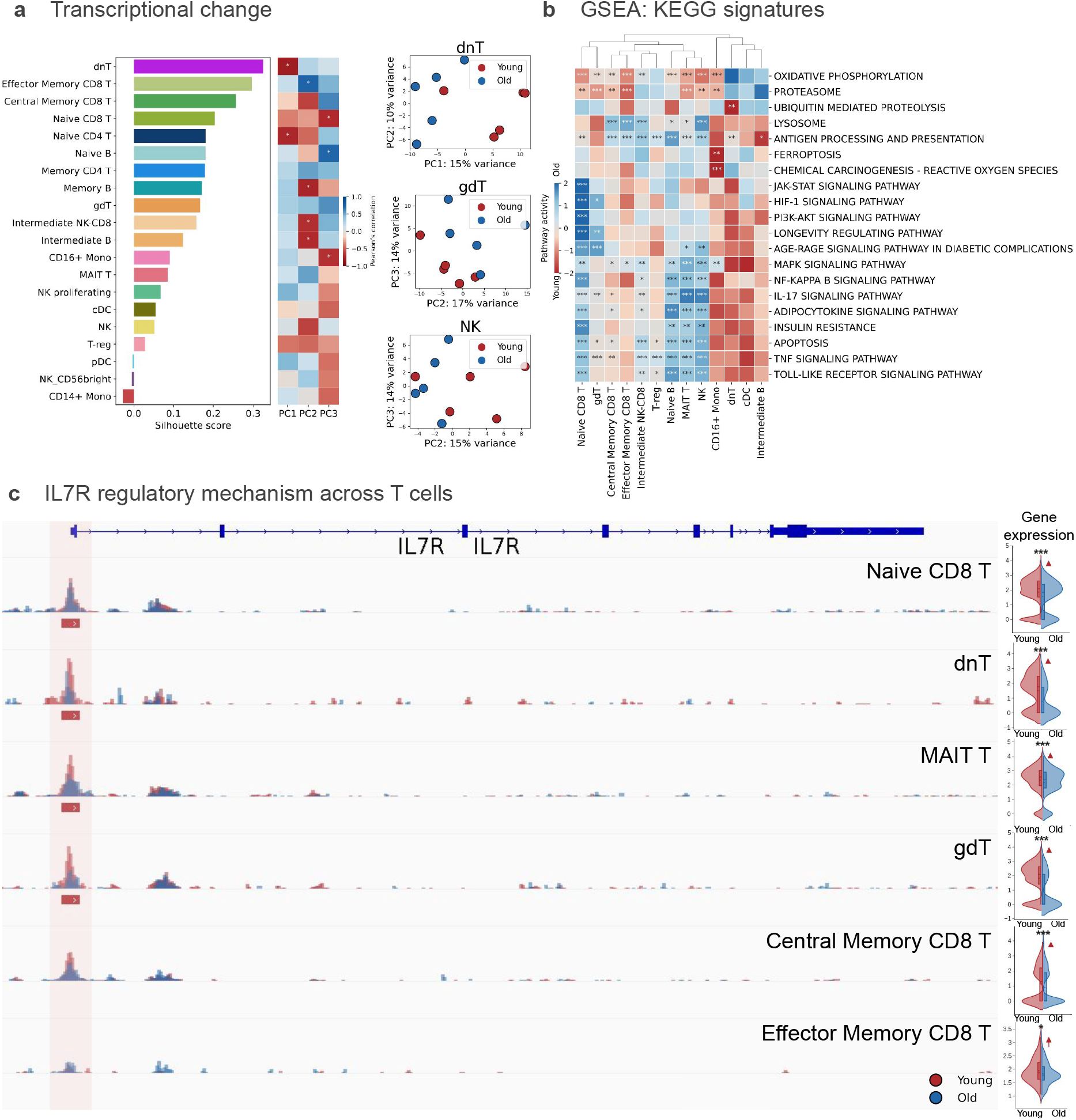
a) Silhouette score reflecting the amount of separability of the cell type specific pseudo-bulk libraries in the first 3 PCs components (left). Cell types are ordered by increasing score, while the highest scores indicate the highest separability. On the right, we showed two components for selected cell types with high separation between old and young donors. b) KEGG pathway enrichment in differential expression genes significant in at least one cell type. The two-sided p-values derived from the t-test of the regression coefficient in the univariate linear model as ^*\*\*\**^*p* < 0.001, ^*\*\**^*p* < 0.01, ^*\**^*p* < 0.05. c) Differential accessibility between young and old across T cells (left) along the IL7R expression changes (right) with the two-sided p-values derived from the Mann-Whitney U test as ^*\*\*\**^*p* < 0.001, ^*\*\**^*p* < 0.01, ^*\**^*p* < 0.05, where red color represents young and blue for old groups. Red bars indicate differential peaks found in young donors.

**Supplementary Figure 3:**
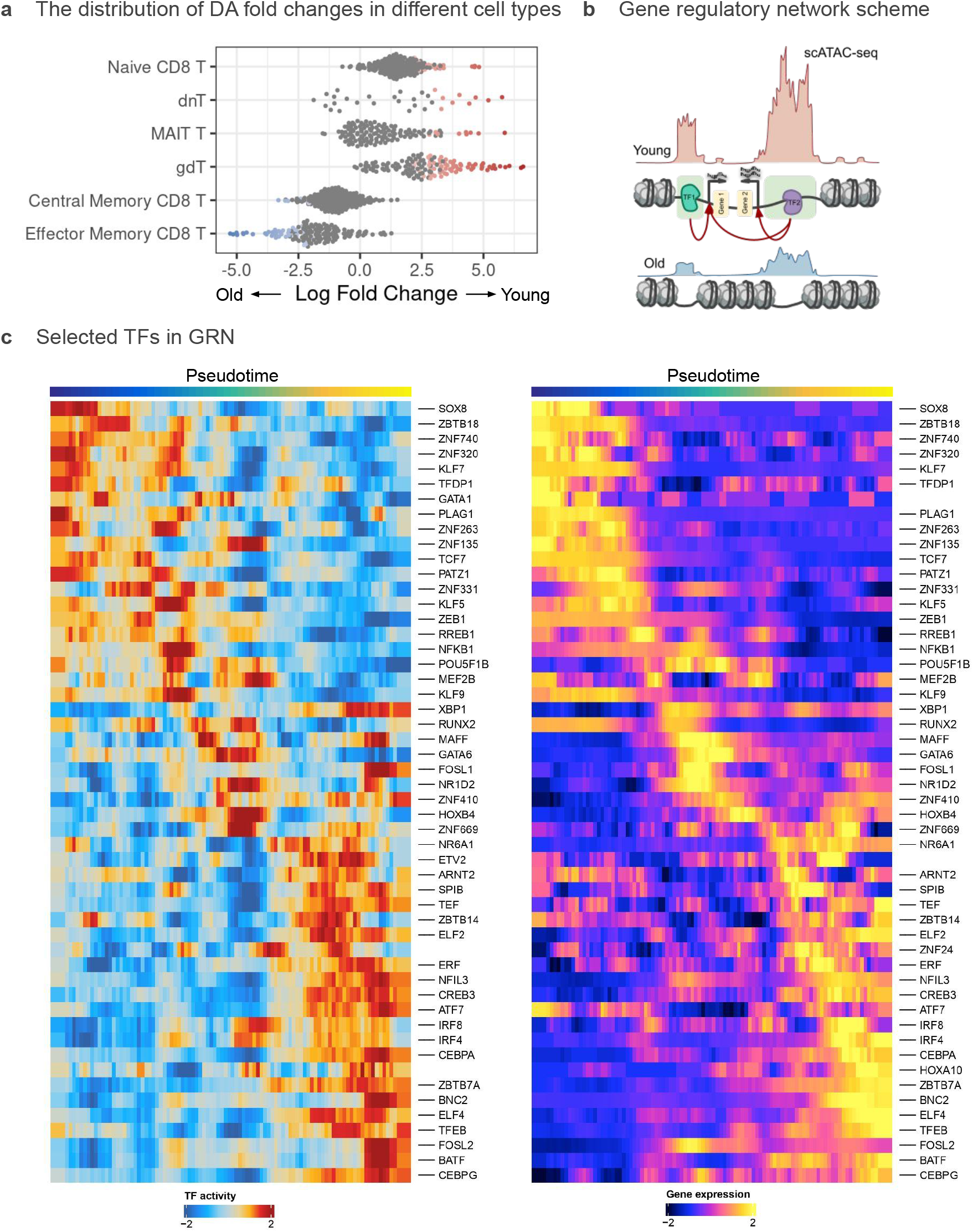
Gene regulatory network in aging. a) Beeswarm plot of differential abundance neighborhoods from KNN graph resulted from Milo. The points are colored by whether they are significant based on the spatial false discovery rate threshold of 0.1. b) Scheme of peak-to-gene link used in gene regulatory network. c) Selecting TFs for GRN which have the highest Pearson correlation (R > 0.4; p-value < 0.01) between TF activities and expressions across the pseudotime trajectory.

**Supplementary Figure 4:**
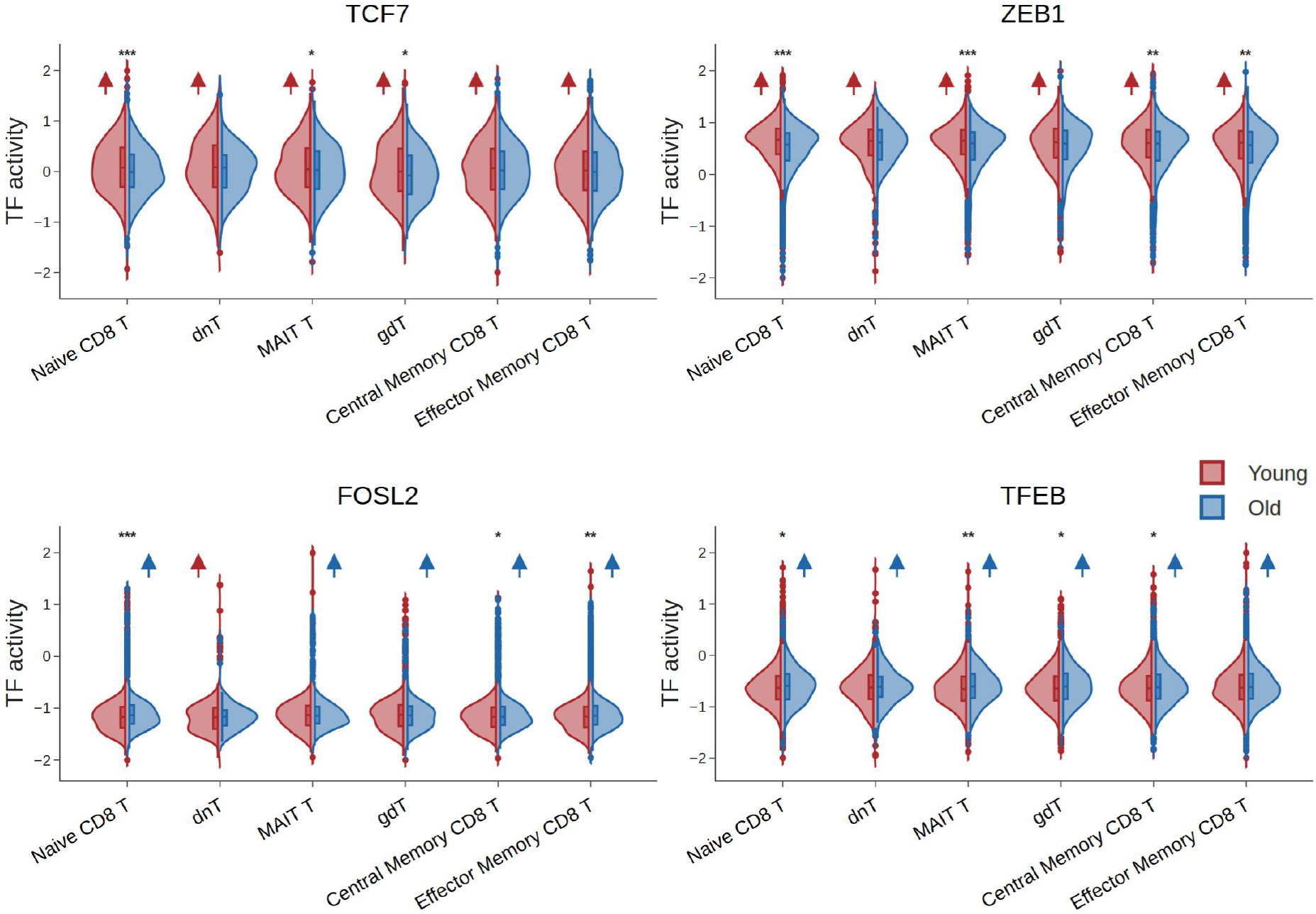
Transcription factor activity changes in aging across pseudotime cell types. While TCF7, ZEB1 show significantly higher activity in young samples, mostly in naïve CD8^+^ T and MAIT T, compared to old samples, the activity of FOSL2, and TFEB is significantly higher in old samples, specifically in Effector Memory CD8^+^ T with the two-sided p-values derived from the Mann-Whitney U test as ^*\*\*\**^*p* < 0.001, ^*\*\**^*p* < 0.01, ^*\**^*p* < 0.05 and an arrow to show the direction of the changes as blue arrow shows higher activity in old samples and red arrow shows higher activities in young samples.

**Supplementary Figure 5:**
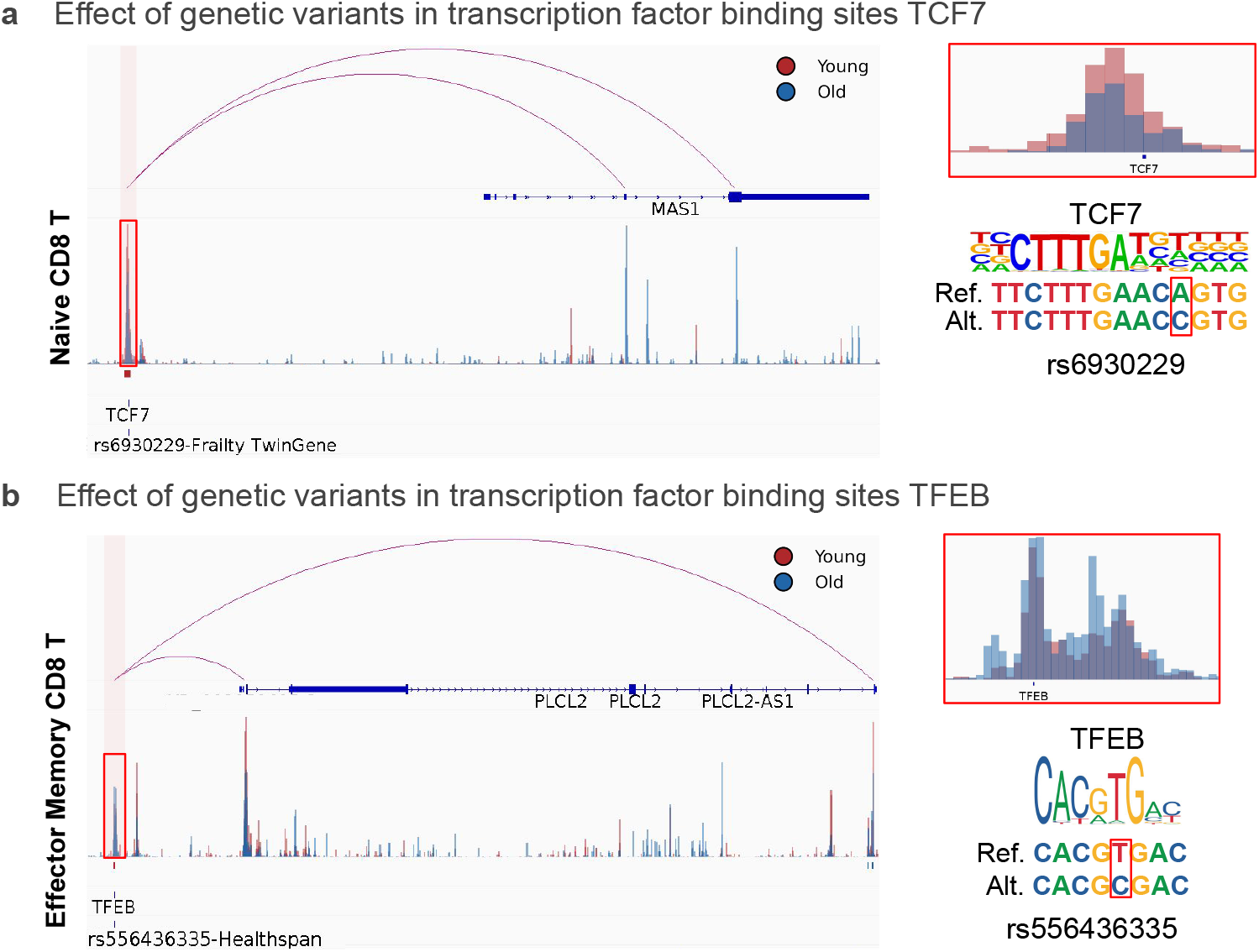
Genetic variants effects in aging. a, b) Effect of genetic variant in transcription factors binding sites of TCF7 and TFEB potentially affecting the regulation of MAS1 and PLCL2 genes, respectively. For the peak close to MAS1, it loses accessibility with age in naïve CD8^+^ T cells, while for the peak close to PLCL2 we observe a gain in accessibility in age in effector memory CD8^+^ T. MAS1 has been shown to be related to aging in model organisms [61]. PLCL2 is a risk factor for autoimmune diseases. [62]

